# Epitranscriptomic Dysregulation Underpins Artemisinin Mechanism of Action

**DOI:** 10.1101/2025.08.28.672746

**Authors:** Ameya Sinha, Benjamin Sian Teck Lee, Junmei Samantha Kwah, Jiaqi Liang, Hannah Murray, Radoslaw Igor Omelianczyk, Sebastian Baumgarten, Peter C. Dedon, Peter R. Preiser

## Abstract

Artemisinin has long been a first-line antimalarial. Yet, its mode of action is still poorly understood. Emergence of artemisinin-resistant strains highlight the importance of addressing this question so as to develop better drugs and overcome resistance. In this study, we performed RNA-sequencing and proteomics studies on artemisinin treated parasites indicated a striking difference in the codon-usage pattern of differentially translated genes. Using a liquid chromatography-coupled mass spectrometry (LC-MS)-based platform, we have quantified the full spectrum of modified ribonucleosides on tRNA in *P. falciparum* in response to the drug. We found that N^6^-threonyl-carbomyladenosine (t^6^A), a universal tRNA modification found at position 37 is hypomodified in response to artemisinin induced stress. Additionally, we also found that artemisinin treatment resulted in a downregulation of PfSua5, an enzyme involved in the t^6^A biosynthesis machinery. These findings provide new insights into how artemisinin works. More broadly, the findings exposes the tRNA epitranscriptome as a vulnerability in the parasite that can be exploited for new drugs.

## Introduction

Malaria is a major life-threatening infectious disease caused by the unicellular, apicomplexan parasite species belonging to the *Plasmodium* genus. It is transmitted to humans through the bite of an infected female *Anopheles* mosquito and persists as a major public health burden today, particularly in tropical and sub-tropical low-income regions of the world with 263 million cases that resulted in 597,000 deaths in 2023^1^. Most of these deaths are attributed to *Plasmodium falciparum*. Despite a drastic reduction in malaria cases and deaths in the past decade, progress has been hampered by the widespread emergence of drug resistance in both parasite^2–8^ and vector species^9, 10^

Understanding and solving the problem of drug resistance requires a deep understanding of how specific drugs interact with the target pathogen, both in the context of their mode of action but also in relation to protective stress responses. Artemisinin is a first-line antimalarial drug that traces its origins to ancient China where the plant *Artemisia annua* was used for various ailments since as early as 168 B.C^11^. The identification of artemisinin as the active ingredient responsible for its antimalarial properties occurred in 1972, with the compound and its derivatives remaining the mainstay of treatment for malaria to this day, where it is often used in combination with other drugs in the form of one of many artemisinin-based combination therapies (ACTs)^12^.

Despite the long history of artemisinin use, the molecular details of its mode of action, stress responses and resistance mechanisms have only begun to surface in recent years. Current thinking proposes that the activation of the drug by cleaving its endoperoxide bridge though a reaction with either haem or free ferrous iron to forms secondary carbon-centered radicals cause widespread damage to proteins^13, 14^. Initial studies showed that these radicals form covalent adducts with several proteins, including the *P. falciparum* Translationally Controlled Tumor Protein homolog (PfTCTP), as well as the SERCA orthologue (PfATP6)^14, 15^. Subsequent proteomic studies have further extended our understanding of the specific targets of artemisinin. A membrane proteomics study found that treatment with the artemisinin derivative dihydroartemisinin (DHA) downregulated *P. falciparum* erythrocyte membrane protein-1 (pfEMP1)^16^, while a chemoproteomic study relying on clickable artemisinin-based activity probes found 124 covalent binding protein targets^17^. All the studies to date suggest that while the activation of artemisinin can cause protein damage this effect is potentially more specific than initially believed. Moreover, recent work profiling the full parasite proteome alongside epitranscriptomic changes in response to artemisinin challenge has shown that protective stress responses involving RNA modifications are linked to resistance of the ring stage parasite to artemisinin^18^, which is particularly relevant to the context of the current work being presented here.

To expand our understanding of the potential mode of action of artemisinin and potential stress response mechanisms we examined the proteome and transcriptomic of *P. falciparum* trophozoites after treatment with DHA. This data showed a discordance in the expression profiles of proteins and mRNA indicating an impact of DHA at the post-transcriptional level and indicated that a shift in codon preferences due to changes in the modification status of tRNAs could explain these observations. Consistent with this model we show that the amount of the t^6^A and the tRNA-modifying enzyme responsible for its installation, PfSua5, were decreased in DHA-treated parasites. Taken together, our findings suggest that epitranscriptomic dysregulation is not only relevant in the context of artemisinin resistance but also in relation to its mode of action. Most importantly this work shows that targeting RNA modifications in *P. falciparum* may represent a new so far underexplored avenue for the development of new therapeutic compounds.

## Results

### DHA exposure results in discordant alterations in mRNA and protein levels

To gain a more comprehensive understanding of the mode of action and associated stress responses to DHA exposure, we performed transcriptomics and proteomics on parasites exposed to IC_20_, IC_50_, and IC_80_ concentrations of DHA for 3 hours at 24hpi. The transcriptomes of the DHA exposed parasites showed 164 mRNAs to be differentially downregulated and 118 to be differentially upregulated at the IC_20_ dosage, 508 differentially downregulated and 524 differentially upregulated at IC_50_, and 787 differentially downregulated and 787 differentially upregulated at IC_80_ (Figure 1a).

**Figure 1:**
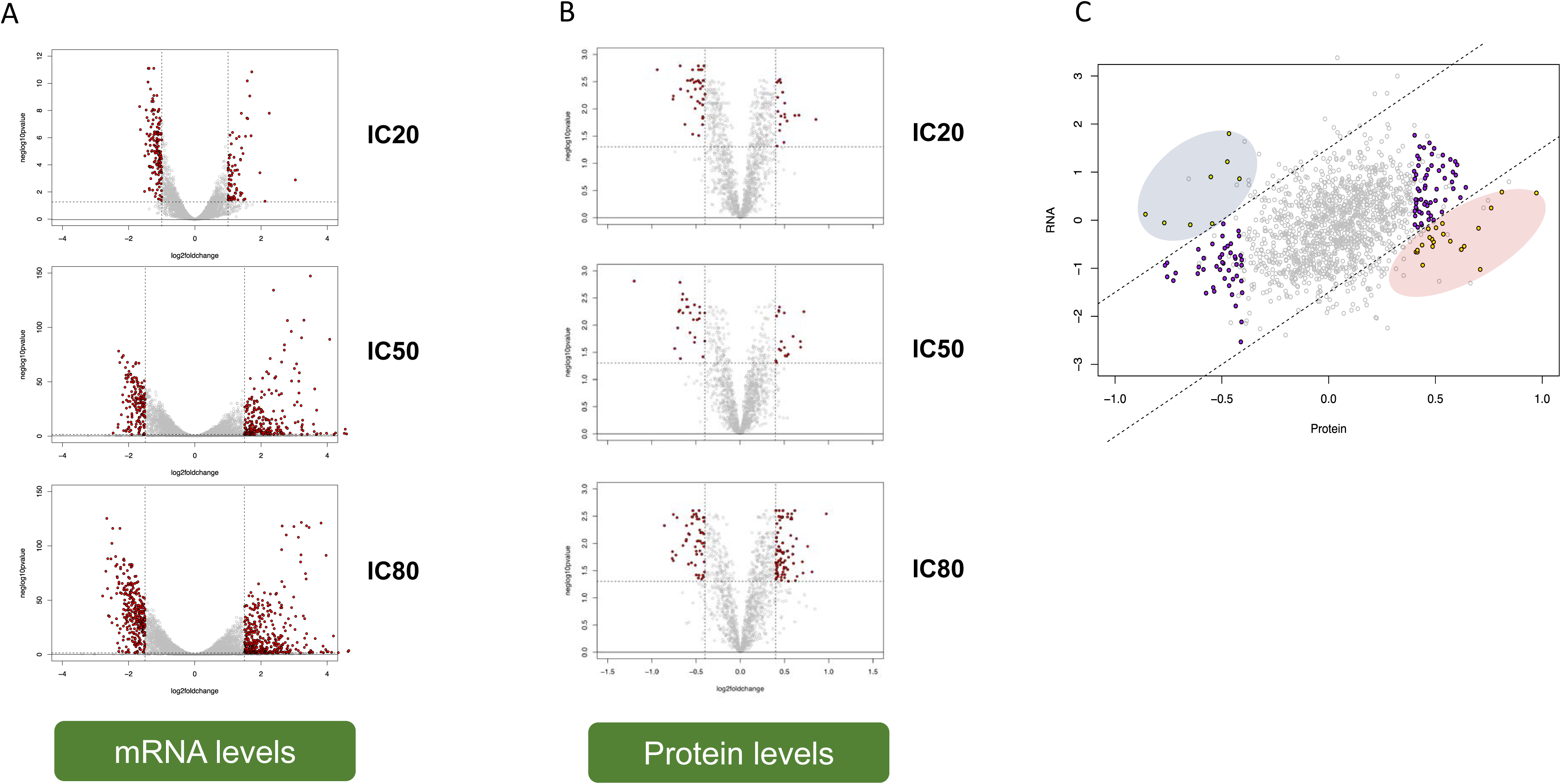
Lack of correlation between mRNA and protein levels upon DHA treatment A. The transcriptomes of the DHA exposed parasites show 164 mRNA to be differentially downregulated and 118 to be differentially upregulated at the IC_20_ dosage, 508 differentially downregulated and 524 differentially upregulated at IC_50_, and 787 differentially downregulated and 787 differentially upregulated at IC_80_, B. The proteomes of the DHA exposed parasites show 49 proteins to be differentially downregulated and 18 to be differentially upregulated at the IC_20_ dosage, 37 differentially downregulated and 13 differentially upregulated at IC_50_, and 55 differentially downregulated and 40 differentially upregulated at IC_80_, C. Comparison of protein and transcript fold changes under DHA exposure at IC80 concentration reveals 8 LTE (blue) and 21 HTE (red) genes.

Using TMT-based quantitative proteomics, which resulted in the coverage of 25.15% of the parasite genome, with a total of 41255 peptide spectral matches (PSMs), and 1243 proteins quantified with a good distribution across all major gene ontology categories, we found the impact of DHA exposure on the proteome of the parasites to be far more muted. At IC_20,_ only 49 proteins were differentially downregulated and 18 differentially upregulated. 37 were differentially downregulated and 13 differentially upregulated at IC_50_, and 55 were differentially downregulated and 40 differentially upregulated at IC_80_ (Figure 1b).

To better understand the relationship between transcriptional and proteomic changes we next compared the Log2FoldChange values of mRNA for each gene to its protein Log2FoldChange. Under DHA treatment at IC_80_, 8 genes were found to have low translational efficiency (LTE), being significantly decreased at the protein level despite no change or an increase at the mRNA level. Conversely, 21 genes were found to display high translational efficiency (HTE) with significantly higher protein abundances despite no changes or a decrease at the mRNA level upon DHA exposure at IC_80_ (Figure 1c). This suggested that post-transcriptional regulation plays a role in the parasites response to DHA.

### DHA challenge leads to the specific reduction of t^6^A

Considering the previous studies that showed that tRNA modifications are important translational regulators during the parasite life cycle^19^ and have been implicated in DHA ring stage resistance^18^, we performed ribonucleoside LC-MS/MS to examine whether changes to the tRNA epitranscriptome could explain the differences in translational efficiency. Small RNAs were isolated from parasites exposed to IC_20_, IC_50_, and IC_80_ concentrations of DHA for 3 hours at 24hpi. An enzyme cocktail was then used to digest the tRNA to its individual mononucleosides which were subsequently subjected to LC-MS/MS.

Among the 30 tRNA modifications that we detected and quantified with this approach, N^6^-threonylcarbomyladenosine (t^6^A) showed the greatest degree of reduction in the treated parasites, with a mean fold change at IC_20_, IC_50_, and IC_80_ of 0.793, 0.732, and 0.769 with an SD of 23.7%, 20.9%, and 20.8% respectively (Figure 2a). m^6,2^A at IC_80_ with a fold change of 1.292 and an SD of 38.5%. This is an rRNA modification and its presence in the small RNA fraction where tRNA is found likely points to the occurrence of rRNA degradation.

**Figure 2:**
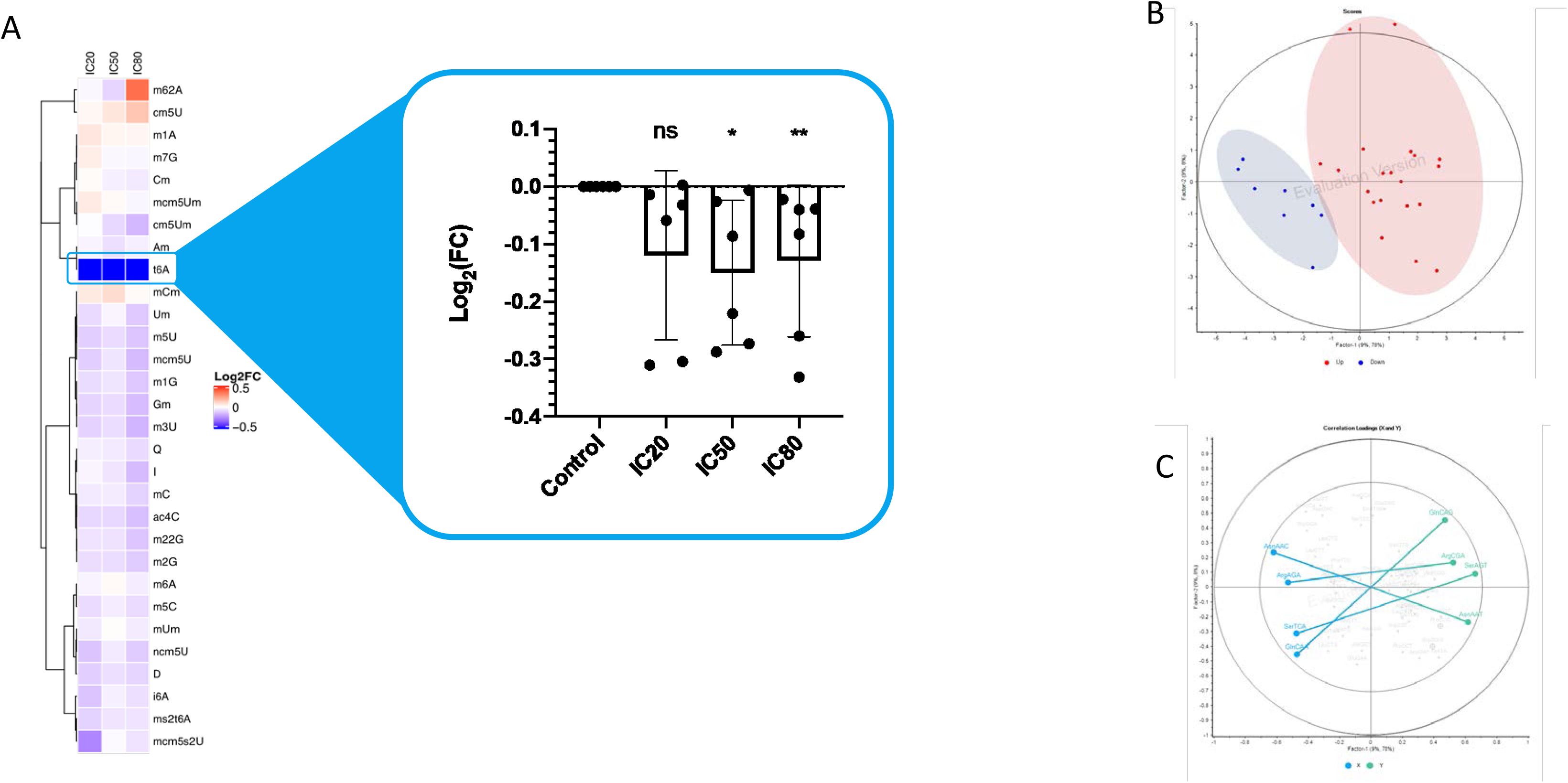
tRNA modification t6A is downregulated upon DHA exposure A. Heatmap of modified ribonucleoside levels show t6A significantly downregulated at IC50 and IC80 DHA treatments, P < 0.05, B. PCA of codon usage percentages of LTE and HTE genes show that these genes are distinguished from each other. The ellipses represents the Hoteling T2 limit, P < 0.01 (F-test), C. The corresponding x, y correlation loadings plot for the analysis shows the codons most strongly contributing to this separation between LTE and HTE genes, with circled codons significantly contributing as determined by PLS analysis. The outer and inner ellipses indicate 100% and 50% explained variance, respectively.

t^6^A is a tRNA modification found on ANN-decoding tRNAs. It is essential and prevents amino acid misincorporation and its downregulation would be expected to disproportionately impact the translation of genes enriched in ANN-codons. To explore this further we investigated the codon preferences of the genes exhibiting LTE or HTE tendencies. PCA analysis clearly distinguished the LTE and HTE genes based on their codon usage preferences (Figure 2b) and more detailed analysis showed that this separation is mainly driven by ANN codons such as AsnAAC, AsnAAT, ArgAGA, and SerAGT (Figure 2c). Taken together this data indicates that the reduction in t^6^A levels in response to DHA treatment contribute to the specific deregulation of translation observed.

### Reduction in t6A levels does not protect against DHA challenge

Reduction of mcm^5^s^2^U in response to artemisinin has been shown to protect ring stage parasites against the drug and we therefore explored whether a reduction of t^6^A was associated with some protection against the drug in trophozoite parasites. To investigate this we first generated a conditional knockdown (cKD) parasite line for PfSua5 (1203), the enzyme responsible for synthesis of the TC-AMP precursor from threonine, bicarbonate, and ATP (Figure 3a). The cKD line makes use of the TetR-DOZI system (Figure 3b) and generates an HA-tagged PfSua5 suitable for subsequent downstream analysis. This system regulates translation at the post-

**Figure 3:**
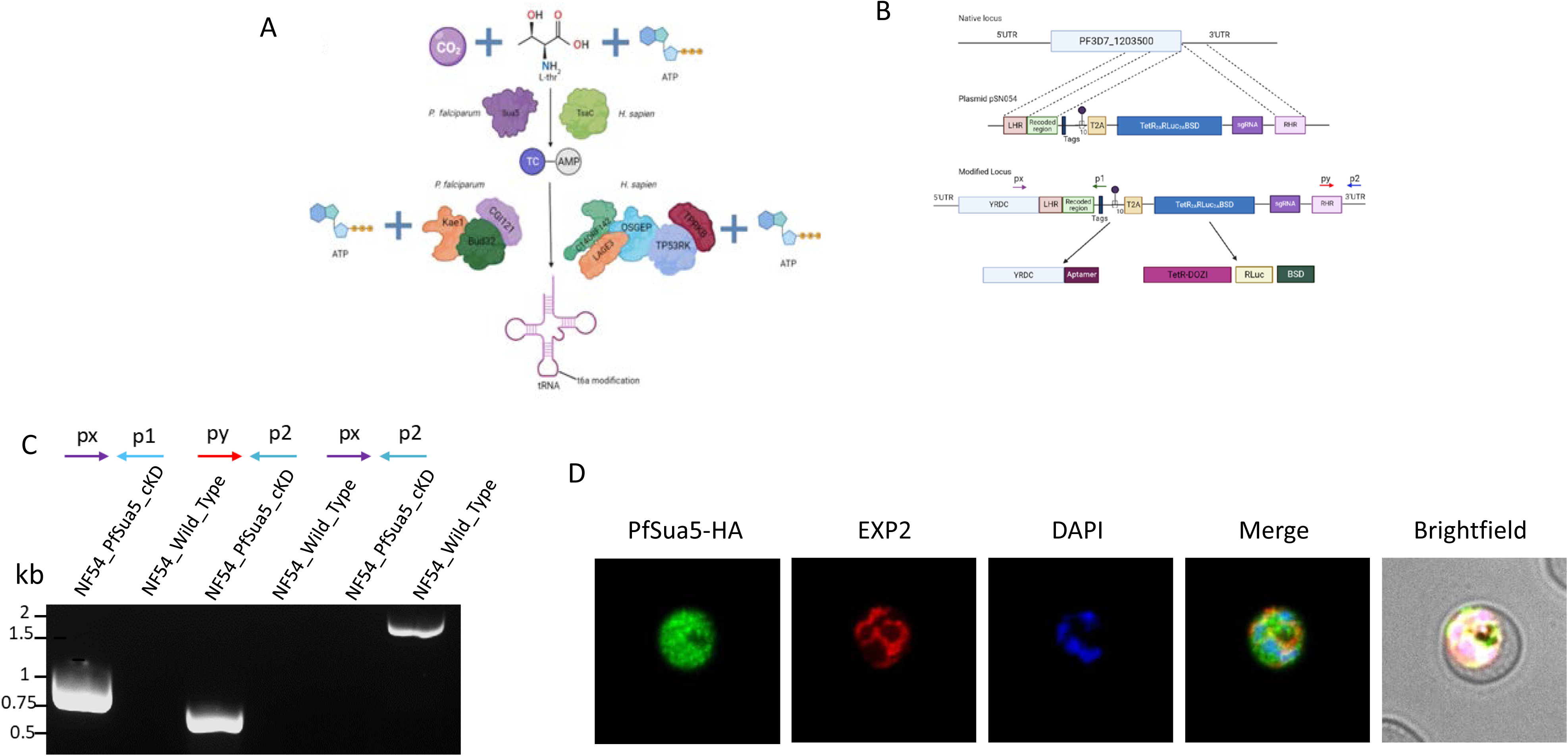
Generation of inducible t6A knockdown line A. Schematic diagram illustrating the biosynthesis of t^6^A in P. falciparum and H. sapiens. In P. falciparum, PfSua5 first synthesises the TC-AMP intermediate from carbon dioxide, threonine, and ATP. An enzyme complex comprising PfBud32, PfKae1, and PfCGI121 then adds this to position 37 on ANN-decoding tRNAs, B. Generation of a parasite line for inducible knockdown of PfSua5. The genomic locus containing the PfSua5 gene is modified to attach a HA-tag for visualization and a TetR-DOZI aptamer to allow its expression to be controlled by aTc. Integration PCR is used to confirm successful chromosomal integration, C. Integration PCR shows successful integration of the cassette into the genomic locus, D. Immunofluorescence assay shows PfSua5 is localized to the cytosol of the parasite.

transcriptional level with the help of anhydrotetracycline (aTc). In the absence of aTc, the mRNA is sequestered into stress granules resulting in a knockdown.

Integration PCR showed successful integration of the HA-tagged, aTc-controlled cassette into the PfSua5 genomic locus (Figure 3c) and immunofluorescence assay showed that PfSua5-HA was expressed and localizes to the cytoplasm of the parasite, remaining within the parasite boundaries stained by the EXP2 antibody and not exiting the parasite to enter the cytosol of the infected RBC (Figure 3d).

To reduce the levels of t^6^A the cKD parasites were cultured for varying duration in the presence and absence of aTc. Western blot analysis showed a significant reduction (68%) of PfSua5-HA after culturing for for half a cycle (24hrs) in the absence of aTc with a further reduction being observed if the duration is increased to 1.5 cycles (5%) (Figure 4a, b). Importantly, the reduction of PfSua5-HA due to aTc depletion resulted in a concurrent reduction of t^6^A abundance (Figure 4c).

**Figure 4:**
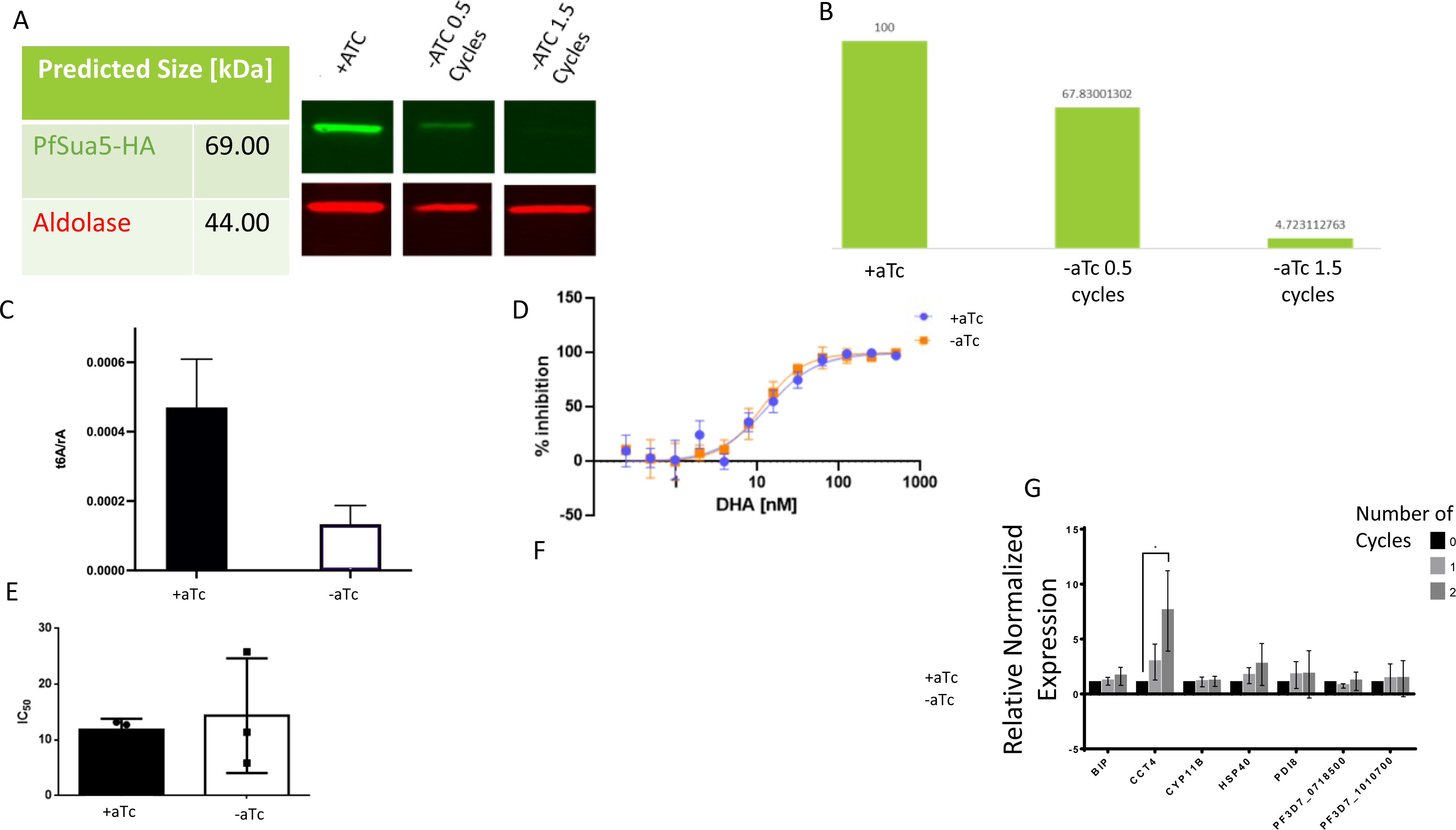
Depletion of PF3D7_1203500 leads to reduce t6A and impacts parasite growth A. Knockdown is observed after removal of aTc for as little as 0.5 cycles, B. Densitometry analysis of the western blot shows that at 0.5 cycles, the expression of PfSua5-HA is 68% of the + aTc condition and at 1.5 cycles it is 5%, C. Ribonucleoside LC-MS/MS shows that parasites cultured in the absence of aTc have reduced levels of t^6^A, D. Dose-response profile of cKD parasite line treated with DHA in +aTc and –aTc conditions are highly similar, E. IC50 value of reveals no significant change in sensitivity of the cKD parasite line to DHA in +aTc and –aTc conditions, F. Growth curve of parasite measured over 9 cycles cultured in the presence (+aTc) or absence (-aTc) of aTc, G. RT-qPCR of UPR genes show significant upregulation of CCT4 mRNA after two cycles in the absence of aTc, P < 0.05.

**Figure 5:**
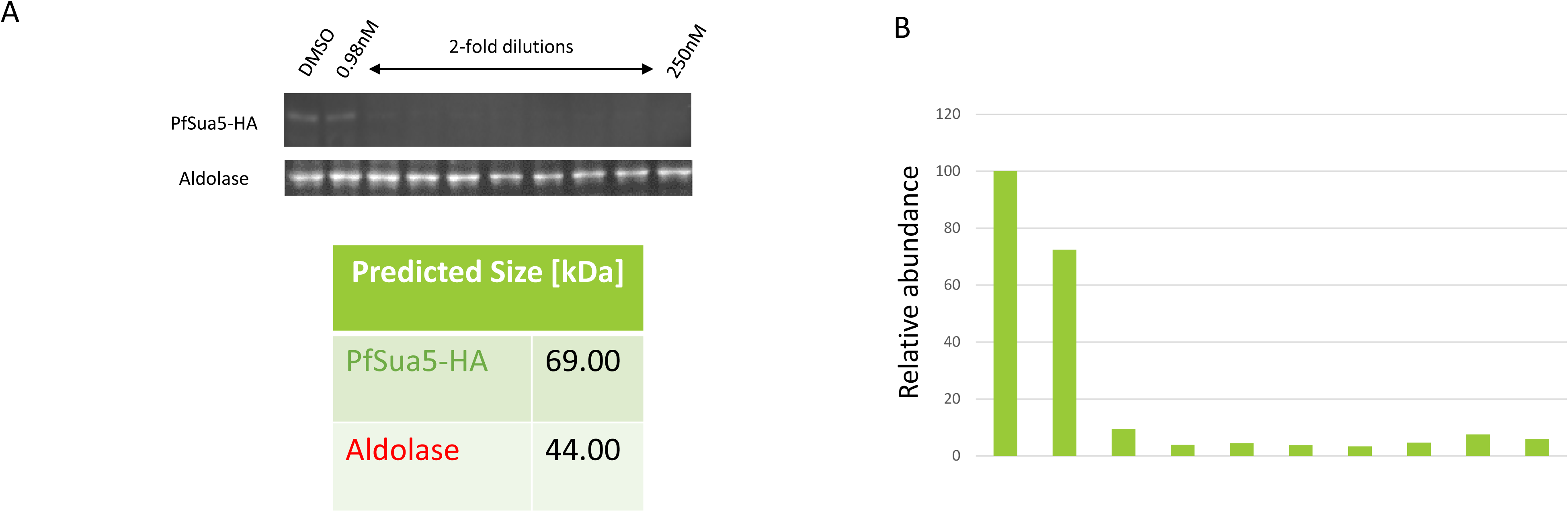
Exposure to DHA reduces PfSua5-HA A. Western blot of PfSua5-HA tagged parasites treated with a 3hr pulse of DHA at varying concentrations show reduction of PfSua5-HA from the 1.9nM dose onwards, B. Densitometry analysis of the western blot shows that the reduced expression is about 80% of the DMSO-treated control.

To explore whether reduction in t^6^A abundance was associated with reduced sensitivity of the parasite to DHA, parasites were cultured for 1.5 cycles in the absence of aTc and the IC50 was determined. These results showed that the reduction of t^6^A abundance does not have a protective effect against the drug (Figure 4d, e).

### Reduction in t^6^A abundance impacts parasite viability

Having demonstrated that a decrease in t6A has no protective effect against DHA we explored whether it impacts parasite viability. For this, we proceeded to measure the growth of the parasites over time in cultures with and without aTc. This shows clearly that the knock down of PfSua5 impedes parasite growth with the parasites showing a significantly reduced growth (Figure 4f).

Decrease in t6A levels in other systems has been linked to protein mistranslation and the activation of unfolded protein response (UPR)^20^ and we therefore explored whether this is also observed in the case of *P. falciparum* particularly as activation of UPR in response to DHA stress has been reported^21–23^. For this, we performed qPCR on several UPR genes^24^. We observed one gene, the Chaperonin Containing TCP1 Subunit 4 (CCT4) to be significantly upregulated two cycles after removal of aTc (Figure 4g), mirroring an observation made during DHA treatment.

### DHA exposure directly reduces PfSua5 abundance

Our data showed that while the reduction in PfSua5 did not provide any protection against DHA it had a direct impact on parasite growth. This suggests that DHA might specifically target PfSua5 protein levels and this contributes to the overall mechanism of parasite killing. To evaluate this PfSua5-HA parasites grown in the presence of aTc were exposed to varying concentrations of DHA. Western blot analysis of parasite extracts prepared after 3 hrs DHA exposures showed a clear impact of the drug on PfSua5 levels with as little as 1.9nM of DHA reducing the protein to nearly undetectable levels. We observed that at DHA concentrations of 1.9nM and above, PfSua5-HA expression drops to fixed level and does not drop further (Figure 4a). Densitometry analysis estimates this level of expression to be around 5% that of the DMSO-treated control (Figure 4b). This is consistent with our LC-MS/MS data that showed that IC_20_ DHA concentrations already led to a maximum reduction of t^6^A.

## Discussion

Although artemisinins are first line antimalarials, the emergence of parasite strains that are resistant to it poses a threat to its utility^2–8^. Overcoming this resistance demands that we understand its mechanism of action better. In this study, we use the artemisinin analog DHA to provide evidence showing that the tRNA-modifying enzyme PfSua5 could be a biologically relevant target of artemisinin.

PfSua5 catalyzes the formation of t^6^A, one of several complex modifications to be found on tRNA. This modification occurs at position 37 of ANN-decoding tRNA isoacceptors^25^, where it stabilizes the pairing between A_1_ of the codon and the U_36_ of the anticodon to prevent frameshifting during translation^26^, while facilitating translocation of the tRNA from the aminoacyl site to the peptidyl site in the ribosome^27^.

The greater rate of frameshifting that is expected to accompany the loss of t^6^A could explain the proteotoxic stress observed in DHA-treated parasites^22, 23, 28^. This finding also aligns well with evidence in the literature showing that inhibiting protein translation with the drug cycloheximide antagonizes DHA-mediated killing^22^, since doing so would limit the amount of damage done by aberrant tRNA isoacceptors that lack t^6^A. Implication of the ubiquitin-proteasome system (UPS) in artemisinin mode of action^22, 23^ also lends support to this line of reasoning as the proteasome would be responsible for getting rid of the mistranslated proteins and disruption of its function would also result in accumulation of this damage.

The link between DHA treatment and t^6^A downregulation becomes even more interesting when considering that the UPR has been found to be associated with artemisinin resistance^24^ and that it is also activated by DHA^22, 23^. Here, we report that, like in other organisms, downregulating t^6^A alone through knockdown of its writer is sufficient to trigger the UPR, potentially explaining why UPR activation occurs during DHA treatment in the first place. The association between the UPR and artemisinin resistance suggests that this sequence of events, where DHA interferes with t^6^A formation and subsequently causes UPR activation, is a clinically relevant one.

Besides contributing to the understanding how artemisinin affects the parasite, our study also provides evidence to support the targeting of the epitranscriptome in the treatment of malaria. Modifications to RNA molecules demonstrably affect gene expression by affecting RNA stability^29–35^, RNA structure and folding^36–38^, localization^39, 40^, and protein translation^33, 41, 42^. Among the classes of RNA, tRNA carries the greatest diversity of modifications. Being central to the protein synthesis machinery, the modifications on these molecules offer organisms with a way to globally reprogram translation in a rapid manner to cope with environmental stressors^43–47^.

Similar to other organisms, tRNAs in *P. falciparum* are heavily modified with at least 28 different ribonucleoside modifications, including m^6^A (N6-methyladenosine), m^5^C (5-methylcytosine), and more complex modifications such as mcm^5^U (5-methoxycarbonylmethyluridine)^18, 19^. These modifications control biological processes like gametocytogenesis^48^, are developmentally regulated across the intraerythrocytic life cycle, and also modulate the sensitivity of the parasites to various drugs like DHA, atovaquone, and apicoplast inhibitors^18, 48^.

With a paucity of transcription factors^49, 50^, the *P. falciparum* is heavily reliant on post-transcriptional means of regulating gene expression. Proposed mechanisms for post-transcriptional control in *P. falciparum* include mRNA processing and degradation^51–, 56^, translational repression^54, 57–61^, translational regulation by untranslated regions (UTR) in mRNA^62–64^, and endogenous anti-sense transcripts^65–70^. However, these gene-specific mechanisms which have local effects do not explain a more global systematic scale mechanism involved in the control of gene expression at the level of translation, for which tRNA modifications are well suited for the job. Our findings show that these modifications are potentially vulnerabilities that can be exploited for the development of new therapies against malaria.

## Conclusion

The emergence of artemisinin resistant strains of *P. falciparum* highlights an urgent need to be better understand the mode of action of this front-line antimalaria. Through transcriptomic, proteomic, and epitranscriptomic profiling, our study identifies the tRNA-modifying enzyme PfSua5 and its modification t^6^A as potentially targets of DHA. Not only does our study shed light on the molecular activities of this drug, it also exposes a underexploited vulnerability in the biology of the parasite that can be leveraged upon for the development of new therapeutic strategies against malaria.

## Materials and Methods

### Plasmodium falciparum parasite culture

In order to culture the asexual stages of *P. falciparum*, strain 3D7 was obtained from MR4 resources. These were grown in human red blood cells (RBCs) in RPMI 1640 medium (Invitrogen) supplemented with 0.5% Albumax II (Gibco, Life Technologies), 23.81 mM (2 g/L) sodium bicarbonate (Sigma), 0.1 mM hypoxanthine (Sigma), and 104.7 μM (50 mg/L) gentamycin (Gibco, Life Technologies). The cultures were incubated at 37 _C with 5% CO2 and 5% O2. The cells were routinely inspected using microscopy to check for culture contamination. Cells were synchronised by repeated treatment with 5% D-sorbitol (Sigma-Aldrich) every two to three generations to enrich ring-form infected and uninfected RBCs. After each sorbitol treatment, the mixture was centrifuged at 600 X g (acceleration, 9; deceleration, 2) for 5 min to remove sorbitol and lysed infected RBCs in the supernatant. The pelleted RBCs were washed twice with incomplete RPMI 1640. Following this enrichment process, the synchronised cells were grown to reinvade fresh RBCs.

### RNA extraction and digestion

To minimise host RNA contamination, parasites were isolated by lysing the RBCs with 0.05% saponin (Sigma) and washed 3-times with ice-cold phosphate-buffered saline (PBS). Isolated parasites were then homogenised with 5 volumes of Trizol reagent (Invitrogen), followed by a 5-min incubation at ambient temperature before freezing at -80°C. The Trizol homogenised cell lysates were incubated at room temperature (RT) for 3 min with chloroform which was one-fifth the lysate volume. The mixture was centrifuged at 12,000 X g for 15 min at 4°C and the aqueous phase was collected. Absolute ethanol (Merck) was added to the aqueous phase (35 % v/v) and total RNA species were extracted using a PureLink miRNA Isolation Kit (Invitrogen) according to manufacturer’s instructions. A sequential isolation protocol was adopted to enrich the yield of tRNA using 70 % ethanol for further experiments. The quality and quantity of RNA extracted from each experimental time point were determined by Bioanalyser using small RNA chips (Agilent Technologies). Purified *P. falciparum* tRNA was hydrolyzed enzymatically as described with a slightly modified protocol using following components in the buffer mix at final concentrations of 10 mM Tris (pH 7.9), 1 mM MgCl2, 5 units Benzonase (99% Purity Merck), 50 μM Desferroxamine (Sigma), 372.2μM (0.1 μg/μL) Pentostatin (Sigma), 100 μM Butylated hydroxytoluene (Sigma), 2 mM (0.5μg/μL) Tetrahydrouridine (Calbiochem), 5 units Bacterial Alkaline Phosphatase (Thermo Fisher) and 0.05 units Phosphodiesterase I (Affymetrix). The enzymes were cleaned up using a 10 kDa spin filter (Omega Nanosep) and the sample was lyophilized and concentrated in 0.1% FA/H2O for LC-MS analysis.

### Analysis of tRNA modifications by LC-MS/MS

Purified P. falciparum tRNA from three biological replicates of selected time points were hydrolyzed enzymatically as described previously. Hypersil GOLD a Q column (100 x 2.1 mm, 1.9 μm, Thermo Scientific) was used to resolve the digested ribonucleosides in a two-buffer eluent system, with buffer A consisting of water with 0.1% (v/v) formic acid and buffer B consisting of acetonitrile with 0.1% (v/v) formic acid. All solvents used were LC-MS grade. HPLC was performed at a flow rate of 300 μL/m. The gradient of acetonitrile with 0.1% (v/v) formic acid was as follows: 0-12 min, held at 0%; 12-15.3 min, 0%-1%; 15.3-18.7 min, 1%-6%; 18.7-20 min, held at 6%; 20-24 min, 6%-100%; 24-27.3 min, held at 100%; 27.3-28 min, 100%-0%; 28-41 min, 0%. The HPLC column was directly connected to an Agilent 6490 triple quadrupole mass spectrometer (LC-MS/MS) with ESI Jetstream ionization operated in positive ion mode. The voltages and source gas parameters were as follows: gas temperature -50 deg C; gas flow - 11 L/min; nebulizer - 20 psi; sheath gas temperature - 300°C; sheath gas flow - 12L/min; capillary voltage - 1800 V and nozzle voltage - 2000 V.

### DHA treatments for IC_50_ Assay

Inhibitory concentration (IC) assays were performed in 48-well plates with two-fold serial dilutions of dihydroartemisinin (Sigma) at 2.5% hematocrit and 1% parasitemia at the trophozoite stage. These cultures were incubated for 48 h to be allowed to propagate for one life cycle for a lifecycle IC50 assay. For short term exposure, the drug treatment was performed for 3 h at 5% parasitemia. The drugs were washed, and the parasites were diluted with fresh blood to 1% parasitemia and 2.5% hematocrit for 48 h and allowed to propagate. The parasites were exposed to different stress agents for 3 hours for the tRNA modification profiling prior to harvesting as described. The experiments were performed with three biological replicates for individual stress exposure. Parasitemia of treated and untreated cultures were quantified by flow cytometry where Hoechst 33342 (Invitrogen) and MitoTrackr (Invitrogen) were used to stain DNA and mitochondria thus distinguishing live cells from dead parasites. For plotting dose-response curve, the concentrations of the treatments were log-transformed, and the responses were expressed as the fraction (%) of treated to untreated control. Nonlinear regression curve fitting was performed with four-parameter least-squares fitting method by Prism 6.0 (GraphPad). The stress agents’ IC20, IC50, and IC80 are determined from the fitted curves and are listed in the results section.

### Protein extraction and digestion

Parasite infected red blood cells were pelleted at 2200 X g for 4 min at ambient temperature and subsequently lysed with ice-cold 0.15% (w/v) saponin (Sigma). The lysate was thoroughly mixed using an aspiration pipette and incubated on ice for 10 min to ensure complete lysis of the RBC membrane. The parasite pellet was centrifuged at 4000 X g for 10 min at minimum deceleration and washed twice with ice cold PBS and frozen at -80°C. The parasite pellet was resuspended in 6x volume of 8M urea containing 1 mM sodium orthovanadate and homogenized using a sonicator pulse for 3 min at 25% amplitude and 2 sec on, 3 sec off pulse time. The lysate was spun at 16,000 X g at 4 _C for 30 min to pellet the insoluble fraction and the lysate was transferred into a new tube. Protein (100 μg) was reduced with 10 mM DTT at 56°C for 1 h and followed by reduction using 100 mM IAA for 1 hour in the dark. This solution was diluted to 1 M urea and digested with 2 μg Trypsin (Thermo) overnight at ambient temperature. The resulting peptides were desalted using Pierce desalting columns as per manufacturer’s instructions. These peptides were reconstituted in triethylammonium bicarbonate (TEAB) and labelled using tandem-mass-tag TMT labels (Thermo) as per manufacturer’s instructions. The labelled peptides were combined, dried, and reconstituted in 0.1% FA. After checking for labelling efficiency, these peptides were then further fractionated using high-pH fractionation columns (Pierce) as per manufacturer’s instructions into 8 fractions.

### LC-MS/MS analysis for proteomics

Peptides were separated by reverse phase HPLC (Thermo Easy nLC1000) using a precolumn (Thermo) and a self-pack 5 μm tip analytical column (15 cm of 5 μm C18, New Objective) over a 140 min gradient before nanoelectrospray using a QExactive HF-X mass spectrometer (Thermo). The mass spectrometer was operated in a data-dependent mode. The parameters for the full scan MS were: resolution of 70,000 across 350-2000 m/z, AGC 3e^6^, and maximum IT 300 ms. The full MS scan was followed by MS/MS for the top 10 precursor ions in each cycle with a NCE of 28 (34 for TMT samples) and dynamic exclusion of 30s. Raw mass spectral data files (.raw) were searched using Proteome Discoverer (Thermo) and Mascot version 2.4.1 (Matrix Science). Mascot search parameters were: 10 ppm mass tolerance for precursor ions; 15 mmu for fragment ion mass tolerance; 2 missed cleavages of trypsin; fixed modification was carbamidomethylation of cysteine; variable modifications were lysine labeled TMT residues, peptide N-terminal TMT labels, methionine oxidation and serine, threonine and tyrosine phosphorylation. Only peptides with a Mascot score greater than or equal to 25 and an isolation interference less than or equal to 30 were included in the data analysis. For TMT samples, a minimum abundance of 500 ion counts was used as a threshold in order to ensure robustness of data.

### Immunofluorescence assay and microscopy

For immunofluorescence microscopy, infected RBCs were allowed to sediment onto poly-d-lysine coated coverslips for 10 min, followed by fixation in 4% paraformaldehyde for 20 min. Following fixation, coverslips were permeabilised with 0.1% Triton-X for 10 min and then blocked with 3% bovine serum albumin (BSA)-PBS for 30 min. After blocking, the coverslips were probed with primary antibodies in 1% BSA-PBS for 1 h at RT, followed by secondary antibody staining in 1% BSA-PBS for 1 h at RT. The coverslips were mounted in Vectashield supplemented with 4’.6’-diamidino-2-phenylindole (DAPI) for nuclease staining. Fluorescence microscopy was performed on a Leica DM0500MB using a 60X oil immersion objective lens and documented with a Leica DC200 digital camera system.

### Western Blot analysis

Trophozoite-stage parasites were separated from the host red blood cells by incubation with 0.15% saponin lysis buffer on ice. Saponin treated parasite pellets were frozen at -80°C until use. Parasite pellets were resuspended in NuPAGE sample buffer (Thermo Fisher) containing 2% β-mercaptoethanol followed by boiling at 95°C for 5 min. Proteins were resolved by SDS-PAGE on 10% gradient gels and transferred to nitrocellulose membranes. Membranes were blocked using Licor blocking buffer and probed with primary anti-rat and anti rabbit antibodies overnight at 4°. They were incubated for 1h with the IR680 and IR800 conjugated secondary antibodies. Protein bands were detected on the Odyssey infrared imager (LI-COR Biosciences) or ChemiDoc (Bio-Rad). The Western blot images were processed, analyzed and quantified using the Odyssey infrared imaging system software.

### Data processing and statistical analysis

Abundances of RNA modifications were normalized to canonicals rA, rU, rG, and rC to account for total RNA amount injected. These were then transformed to log_2_ ratios of modification levels in each time point/dosage relative to either an arbitrary average/untreated control across all samples, respectively. Unless otherwise stated, all data are represented as mean_SD. Proteomics data for each treatment was analyzed by principal component analysis (PCA) using a non-linear iterative partial least squares (NIPALS) algorithm. For interpretations of the relationships between codon usage (codon frequency) and up-regulated and downregulated proteins at different time points, PLS-R was performed using the NIPALS algorithm. The values of codon usage in synonymous codon choices of those proteins were retrieved from the pre-calculated genome-wide codon usage as provided by the Begley Lab. The fold change values from the proteomics data were used as input for the response variable in the PLS-R analysis. The codon usage was charted out as the predictors. Outliers which could cause overfitting were removed by manual inspection of residual sample variances and leverages, as well as Hotelling’s T2 statistics. Marten’s uncertainty test with optimal number of PCs was used to cross-validate the PLS model. Eigenvector based multivariate statistics was performed using UnscramblerX.

## Acknowledgements

P.R.P., P.C.D., J.L., H.M., R.I.O. and A. S. were supported by the National Research Foundation Singapore under its Singapore-MIT Alliance for Research and Technology (SMART) Centre, Antimicrobial Resistance IRG and Singapore Ministry of Education Academic Research Fund Tier 2 (MOE2018-T2-2-131). A. S. acknowledges support from the Singapore-MIT Alliance (SMA) Graduate Fellowship. Proteomics work was performed in part in the Center for Environmental Health Sciences BioCore, which is supported by Center grant P30-ES002109 from the National Institute of Environmental Health Sciences. We would like to thank Dr. Jacquin Niles for providing us with the NF54 parasites and the Psn054 plasmid. We also acknowledge Peiying Ho and Tan Tse Mien for providing support at SMART laboratories in Singapore.

